# 2D Skeletal Muscle Thin Film Actuators Enhance Efficiency of Biohybrid Robots

**DOI:** 10.64898/2026.05.05.723017

**Authors:** Maheera Bawa, Arielle Berman, Laura Schwendeman, Ferdows Afghah, Seanbiron Johnson, Ritu Raman

## Abstract

Biohybrid robots combining compliant synthetic support structures with biological actuators could enable future applications ranging from precision microsurgery to unmanned exploration. Machines actuated by living skeletal muscles are capable of adaptive behaviors, such as sensing and responding to environmental stimuli in real-time, offering functional advantages over non-biological actuators. However, typical skeletal muscle-powered biohybrid robots depend on 3D tissues which require large cell volumes and offer limited control of muscle fiber alignment, thus reducing efficiency of force generation and transduction. Here, we present a locomotive biohybrid robot powered by 2D monolayers, or thin films, of precisely aligned skeletal muscle fibers on a micropatterned hydrogel skeleton. We demonstrate how varying skeleton design parameters, ranging from material stiffness to microscale topology, impacts muscle fiber alignment and resultant actuation strains, generating forces 10X higher than previous 2D skeletal muscle actuators, improving untethered actuation longevity by ∼4500X from < 10 minutes to > 30 days, and increasing efficiency of muscle force output (force per unit volume of muscle) by 20X as compared to 3D muscles. Utilizing our optimized design for skeletal muscle thin films, we create a multi-limbed robot composed of independent muscle-powered fins capable of on/off control and frequency-dependent speed control. With these control inputs, we achieve steered multi-directional locomotion at speeds up to 4 body lengths per minute in straight movement and 1200 degrees per minute in rotational movement, highlighting potential for such actuators to be transformed into long-lasting functional soft robots.

## INTRODUCTION

Organisms in the natural world can adapt their form and function to meet the needs of an unpredictable environment. Recapitulating these abilities in an engineered system requires using materials that react to their surroundings. Towards this end, force-generating living tissues have been paired with abiotic compliant mechanisms, or skeletons, to form kinetic biohybrid machines. These actuators exploit the advantages of their biotic components, including their mechanical compliance, stimuli-responsiveness, and self-healing ability, to enable next generation applications in medicine and robotics.^[1–4]^

Existing kinetic biohybrid robots are driven by the contraction of either engineered cardiac or skeletal muscle. Cardiac muscle, which undergoes spontaneous and rhythmic contraction that can be paced with external stimulation, has been shown to effectively propel biohybrid machines^[5–7]^. Since cardiac muscle can be readily cultured in 2D monolayers, individual cells can be patterned in precise orientations mimicking native tissue to enable efficient force transduction to underlying skeletons.^[8]^ However, cardiac muscle contraction cannot be turned off, and is not designed to operate at frequencies higher than ∼3 Hz.^[9]^ Moreover, due to mechanoelectrical coupling between adjacent cells, specific regions of a cardiac muscle tissue cannot be selectively recruited to scale force production. As a result, cardiac muscle is not well suited for robotic applications that necessitate scalable spatiotemporal control. On the other hand, skeletal muscles can be precisely stimulated to initiate on/off and local spatial control over a wide range of frequencies (< 1 to > 10 Hz)^[10–12]^. To date, engineered skeletal muscle tissues have largely been fabricated in 3D (i.e. cells embedded inside a bulk matrix) via injection molding or extrusion bioprinting-based techniques,^[13–17]^ and have powered robots for a variety of applications including gripping^[12,18,19]^ and locomotion^[11,20–27]^. These 3D muscles are typically stretched around a compliant structure during differentiation, generating isometric tensile forces that promote uniaxial muscle fiber alignment and force generation. However, precise alignment of single muscle fibers within 3D tissues is difficult to achieve via mechanical stretch alone, thus reducing the efficiency of muscle force generation and transduction to a robotic skeleton.^[28]^

Several studies have demonstrated that more precise patterning of skeletal muscle tissues can be accomplished in 2D, where tuning the microtopography of underlying scaffolds enables single cell-level control of fiber alignment and even multi-directional fiber patterning.^[29,30]^ However, early approaches for fabricating aligned 2D skeletal muscle relied on rigid substrates, thus limiting their lifespan as the tissue would delaminate from the substrate within a few days of culture due to the strong passive tension and active contraction forces generated during differentiation.^[31]^ To combat this challenge, recent studies have demonstrated that coating rigid substrates with ultra-soft hydrogels enables longitudinal culture of 2D skeletal muscle monolayers over several weeks.^[32–36]^ Building on these methods, we have developed a 1-step method for using 3D nano-printed molds to precisely template microscale grooves within hydrogels cast on rigid substrates.^[37]^ Our method, termed STAMP: simple templating of actuators via microtopographical patterning, can precisely pattern multi-oriented µm-scale muscle fibers within cm-scale tissues, yielding controllable multi-degree-of-freedom contraction.

While promising, such fabrication techniques for muscle monolayers still rely on tethering the underlying ultra-soft hydrogel (∼0.2 kPa) to a rigid substrate, such as plastic or glass. Since such hydrogels cannot withstand the large passive tension forces generated by muscles, they quickly undergo rapid and permanent shape change when untethered from underlying rigid substrates.^[37]^ As a result, 2D skeletal muscles fabricated on ultra-soft hydrogels have limited lifespans when untethered (< 10 minutes), reducing relevance for long-lasting biohybrid machines. Moreover, reliance on highly compliant substrates for muscle monolayers reduces the tissue’s active contractile force generation capacity (typically ∼100 µN for a cm-scale tissue).^[35]^

Here, we address these critical challenges limiting the performance and real-world application of 2D skeletal muscle-powered robots. Specifically, we establish how tuning hydrogel stiffness and microtopography impacts the alignment, force generation capacity, and longevity of untethered 2D skeletal muscle thin films (**Figure 1a-b**). Following this parametric study, and corresponding computational simulations, we choose design parameters enabling untethered longitudinal culture of functional 2D skeletal muscle for > 30 days, corresponding to a ∼4500X increase in lifespan. Our optimized 2D actuator demonstrates 20X higher force densities than previous 3D skeletal muscle actuators, showcasing a significant improvement in manufacturing efficiency. We then leverage skeletal muscle thin films to power a swimming biohybrid robot composed of 2 independent “fins”. Following demonstrations of selective stimulation, frequency-dependent response, and exercise training of independent muscle fins, we show how spatiotemporally controlled stimulation of independent fins enables untethered steering and multidirectional navigation with tunable speed. This study demonstrates that enhancing the performance and lifespan of 2D skeletal muscle thin films significantly increases the efficiency of biohybrid machines, broadening future applications of skeletal muscle actuators in robotics.

**Figure 1.**
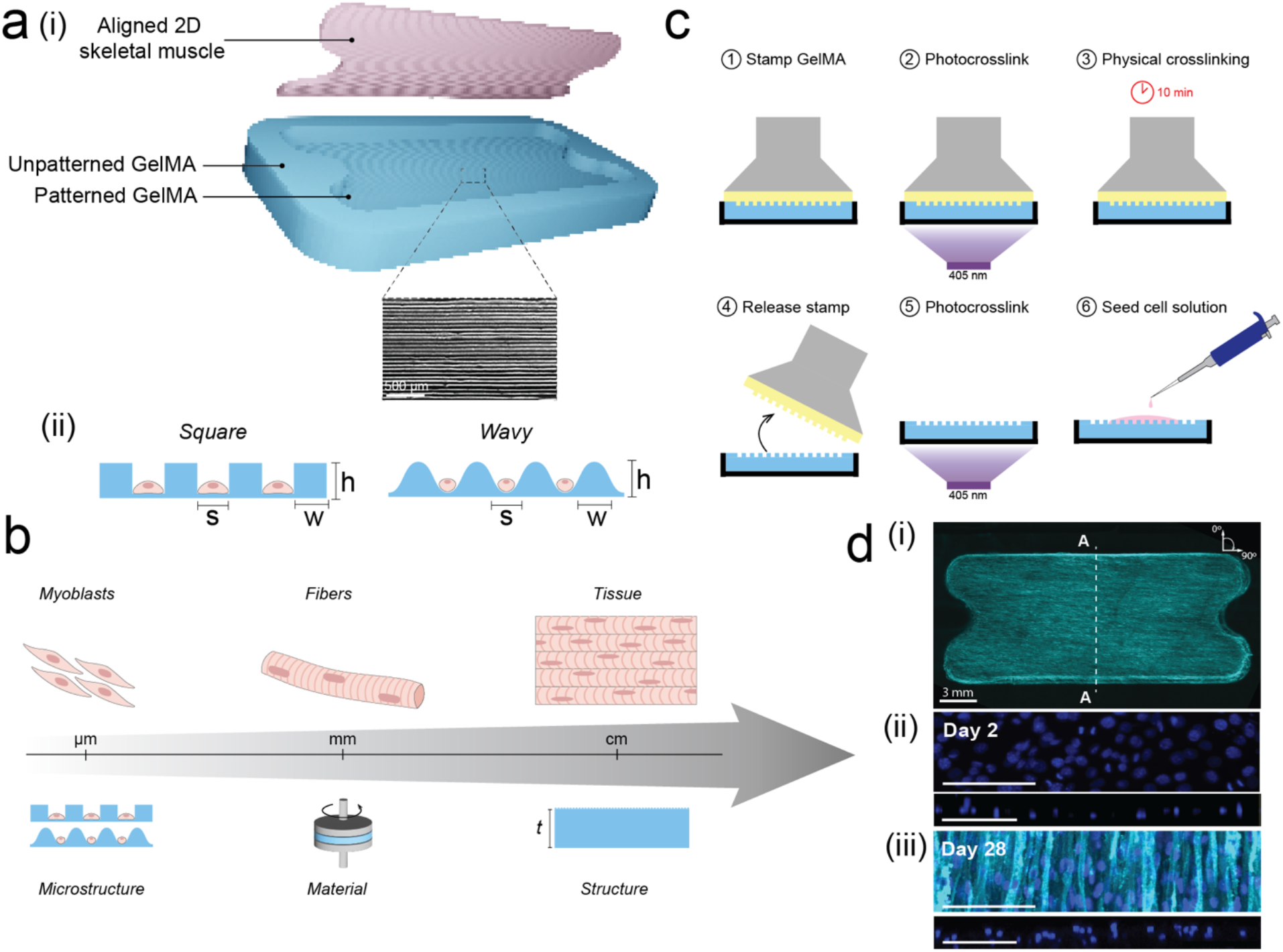
Design and fabrication of 2D skeletal muscle thin film actuators. **(a)** Exploded rendering of the biohybrid actuator with 2D skeletal muscle thin film patterned on an underlying hydrogel (i) with square and wavy microgrooves (ii). **(b)** Hierarchical components of muscle tissue from micrometer-to centimeter-scale and the corresponding skeleton design considerations. **(c)** Fabrication process for the micropatterned hydrogel and subsequent seeding of skeletal muscle cells. **(d)** Representative images of skeletal muscle actuators on hydrogel substrate showing **(i)** top-down fluorescent image of whole tissue with fluorescent membrane tag (TdTomato, cyan) highlighting fiber alignment along 90° grooves. (ii) Zoomed-in top-down and cross-section images of tissues at dashed line labeled “A”, showing nuclei (DAPI, blue) on Day 2 of muscle differentiation. Cells form a monolayer thin film of < 15 µm thickness. (iii) Zoomed-in top-down and cross-section images of tissues at dashed line labeled “A”, showing nuclei proliferation, elongation, and fusion into multinucleated fibers on Day 28 of muscle differentiation. Fibers are aligned parallel to microgrooves in hydrogel scaffold. Scale bars are 100 µm.

## RESULTS

### 2D Skeletal Muscle Thin Film Actuator Design and Fabrication

Reproducible fabrication of 2D skeletal muscle thin films requires a scalable approach for manufacturing cm-scale tissue with µm-scale features. Our previously established STAMP technique enabled templating microtopography into hydrogels cast within standard tissue culture devices (ex: multi-well plates), but did not enable patterning hydrogel substrates of arbitrary shapes that were designed to be untethered. We thus implemented several new STAMP process design features tailored for the fabrication of untethered biohybrid actuators: 1) custom 2-part molds that enabled casting hydrogels in arbitrary shapes; 2) unpatterned raised borders that prevented overgrowth of cells after initial seeding; 3) gas release features within the unpatterned borders designed to limit bubble formation without interfering with desired tissue alignment; 4) a physical curing step for hydrogels to improve stamp release and final micropattern fidelity (**Figure S1**).

We chose gelatin methacrylate (GelMA), a material commonly used in tissue engineering, as the biopolymer substrate for our 2D skeletal muscle substrates for its straightforward compounding and tunable mechanical properties, as well as its established compatibility with 3D skeletal muscle cultures.^[16,38–41]^ Following micro-molding of GelMA scaffolds (7 mm width, 15 mm length, 0.5mm thickness), C2C12 mouse myoblasts were seeded onto the micropatterned area (5.8 mm width, 13 mm length) at a density of 80K cells/mm^2^. Upon reaching confluence in growth media, cells were immersed in differentiation media to promote formation of multinucleated contractile fibers. Following previously established protocols (**Figure 1c**).^[13]^ Our C2C12 muscle line was genetically engineered to express a mutant variant of the 470 nm-responsive calcium ion channel channelrhodopsin-2, enabling precise spatial control of muscle contraction via blue light stimulation of specific tissue regions.^[21]^ Topological patterning of varying cross-sectional geometries (wavy and square) and pitch dimensions (12.5, 25, and 200 µm) in the hydrogel were investigated for their effect on muscle fiber alignment. Following topography optimization, the weight percent of GelMA in the hydrogels (5, 7.5 and 10 wt%) was tuned to elucidate the effect of material stiffness on muscle differentiation and downstream mechanical function.

### Effect of Skeleton Microgroove Design on Actuator Fiber Alignment

Precise alignment of muscle tissues requires understanding the impact of specific microtopologies and dimensions on myoblast fusion and differentiation. We first confirmed that C2C12 cells formed multinucleated fibers on GelMA substrates (**Figure 1d(i)**), yielding a 2D laminar tissue sheet of < 15 µm thickness (**Figure 1d(ii)**).

We then sought to determine the ideal microgroove design by comparing how wavy and square cross-sectional topologies with height/width/spacing of 12.5, 25, and 200 µm impacted the dispersion of the fiber angle from the intended groove angle of 90°. First, holding a constant groove height/width/spacing of 25 µm, we found that square groove architectures yielded a 10X lower average angular dispersion than wavy, flat, and unstamped GelMA samples (**Figure 2a**). This observation may be a consequence of the improved confinement of cells within the square grooves, preventing off-axis fiber growth over the sides of the microgrooves. We then compared square grooves of 12.5, 25, and 200 µm heights/widths/spacings, and found that 25 µm square grooves again resulted in the minimal dispersion from the intended fiber angle (**Figure 2b**). Since 12.5 µm grooves match dimensions comparable to those of single C2C12 myoblasts (∼12 µm in diameter), we hypothesize that natural size variation in myoblasts may prevent adequate confinement in these small groove sizes, and potentially also inhibit fusion for myoblasts that are confined within these grooves. Conversely, 200 µm grooves with features significantly larger than individual cells provided limited control of the directionality of fusion in adjacent myoblasts. Hydrogel skeletons with 25 µm square microgrooves were thus chosen for following characterization studies on 2D skeletal muscle thin film actuators.

**Figure 2.**
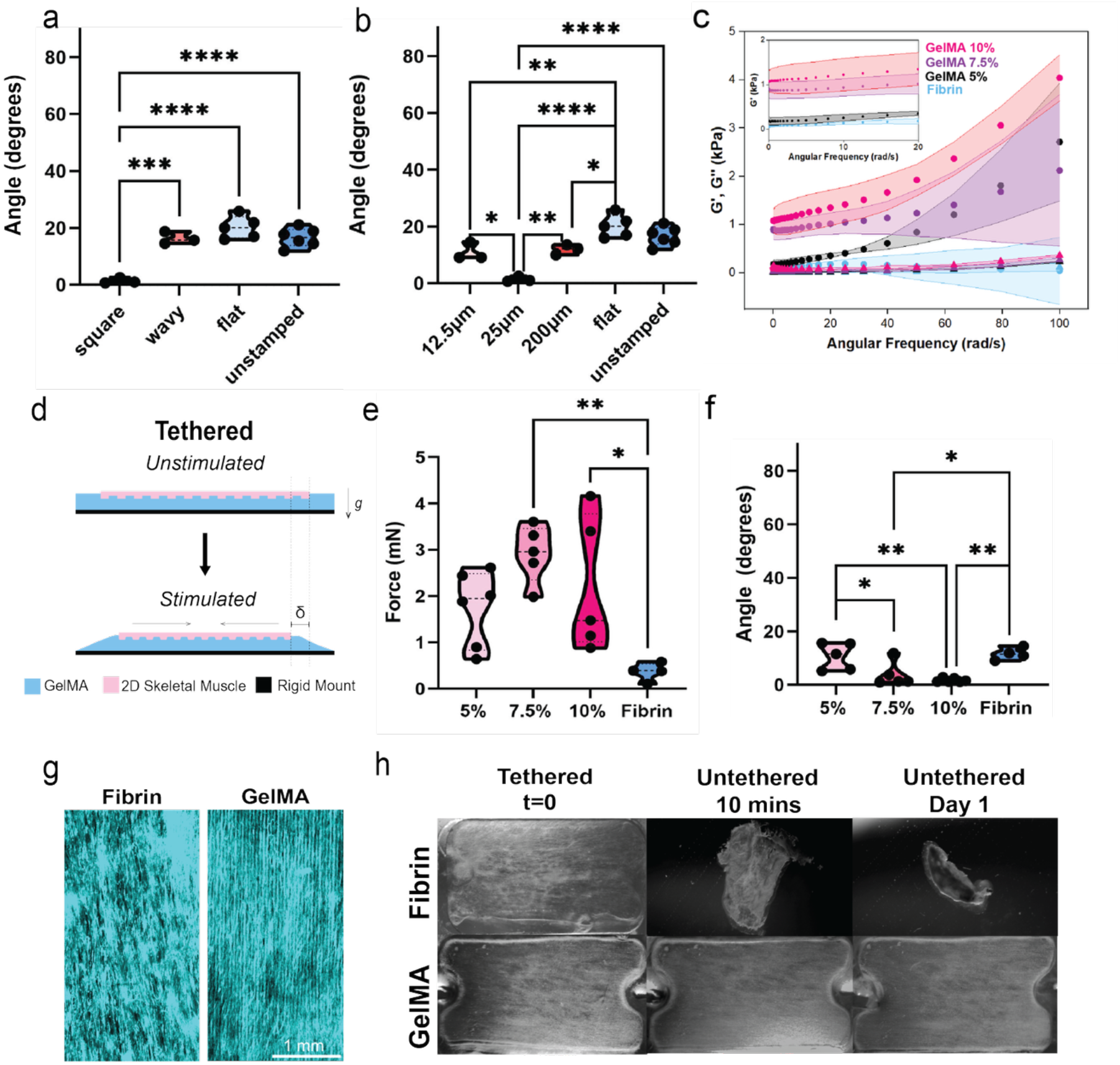
Characterization of hydrogen skeleton parameters on 2D skeletal muscle thin films in tethered and untethered configurations. (a) Dispersion of muscle fibers from desired angle for microgrooves of different shapes compared to a flat stamped controls and unstamped controls. (b) Dispersion of muscle fibers from desired angle for microgrooves of different sizes compared to flat stamped controls and unstamped controls. (c) Rheology results of Fibrin and 5,7.5, and 10 wt% GelMA hydrogels. (d) Tethered muscle actuation schematic. (e) Tethered muscle contractile forces on 5, 7.5, 10 wt% GelMA and Fibrin hydrogels. (f) Dispersion of muscle fibers from desired angle for 25 µm square microgrooves on 5, 7.5, 10 wt% GelMA and Fibrin hydrogels. (g) Fluorescent images of muscle on Fibrin and 7.5 wt% GelMA hydrogel substrates with 25 µm square microgrooves. (h) Time progression of 2D muscle thin films on Fibrin and 7.5% GelMA hydrogels after untethering. In all panels, *, **, ***, and **** labels correspond to p < 0.05, p < 0.01, p < 0.001, and p < 0.0001 respectively.

### Effect of Skeleton Material on Actuator Force and Longevity

Improving the performance and longevity of 2D skeletal muscle thin films requires rheological tuning of the biomaterial substrate to more closely mimic the mechanics of the native extracellular matrix. Optimized matrix viscoelasticity regulates cell differentiation and proliferation and stable tissue-substrate interactions.^[42,43]^ The latter design feature is critical, because the 2D actuators are designed as a bilayer system in which the contraction of the muscle thin film generates tension in the substrate, thus necessitating excellent adhesion between the cells and the hydrogels to prevent delamination of the tissue during long-term untethered actuation.

The stiffness of fibrin hydrogels, the most commonly used extracellular matrix substrates in skeletal muscle tissue engineering, can be tuned by increasing the concentration of fibrinogen monomers in the formulation.^[35,44–46]^ However, the difficulty of dissolving fibrinogen concentrations > 8 mg/mL in cell culture media typically constrains the upper limit of substrate stiffness to < 0.25 kPa.^[35]^ Since GelMA offers a larger range of stiffness tunability^[47]^, we performed rheological characterization of 5, 7.5, and 10 wt% GelMA formulations. At 10 rad/s, corresponding to a 1 Hz actuation frequency that is commonly used to characterize muscle contraction, the storage moduli (G’) of these GelMA formulations were 0.25, 0.94, and 1.2 kPa, respectively, due to their increasing hydrogel crosslinking density (**Figure 2c**). The force output and longevity of biological actuators are both indirectly and directly dependent on these substrate mechanical properties.

### Tethered Contractile Force Output of Skeletal Muscle Thin Films

To precisely quantify how skeleton material properties impacted the magnitude and direction of contractile force generated by 2D skeletal muscle thin films, we first characterized muscle contraction while the actuator was tethered to an underlying rigid substrate (**Figure 2d**). With a known substrate modulus and geometry, the force could be easily calculated using Eq. (1)^[48]^:

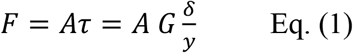

Where *A* is the hydrogel substrate area [mm^2^], *τ* is the shear stress of the substrate caused by the muscle, *G* is the shear modulus of the hydrogel estimated at G’ at 1 Hz [kPa], *δ* is the displacement of the muscle during contraction [mm], and *y* is the thickness of the hydrogel [mm]. Using this calculation, we compared the force output of 8 mg/mL fibrin (G’ = 0.1 kPa at 10 rad/s) to the aforementioned GelMA formulations. We found that 7.5 wt%, and 10 wt% GelMA substrates resulted in the highest muscle thin film forces of 2.9 +/-0.6 mN and 2.2 +/-1.4 mN respectively. By contrast, muscle force output was slightly lower on 5 wt% GelMA (1.7 +/-0.8 mN) and significantly lower on fibrin (0.36 +/-0.19 mN) (**Figure 2e**). We also observed significantly improved fiber alignment in 7.5 wt% and 10 wt% GelMA substrates as compared to both fibrin and 5 wt% GelMA substrates (**Figure 2f**). These experiments highlighted that 7.5 wt% GelMA generated the best aligned and strongest tissues, while using less material than 10 wt% GelMA substrates which produced statistically similar outcomes.

### Untethered Longevity of Skeletal Muscle Thin Films

Beyond force output, longevity of skeletal muscle thin film actuators when untethered is of critical importance for application in biohybrid robots. We fabricated 2D skeletal muscle films on both 8 mg/mL fibrin and 7.5 wt% GelMA substrates containing 25 µm square grooves (**Figure 2g**). We untethered, or fully released the skeleton, from the underlying rigid mount after 2 days of differentiation (**Figure 2h**). The higher G’ of GelMA compared to fibrin prevented the passive tension forces generated by the skeletal muscle monolayer from collapsing the substrate, allowing the GelMA-muscle actuator to maintain its shape (**Figure 2i**) and function for > 30 days (**Figure 3**). On the other hand, the passive tension generated by the 2D muscle tissue caused the ultrasoft fibrin to collapse on itself within 10 minutes of release. Taken together, these results underscore the advantages of 7.5 wt% GelMA as a biomaterial for engineering strong and long-lasting 2D skeletal muscle-powered robots.

**Figure 3.**
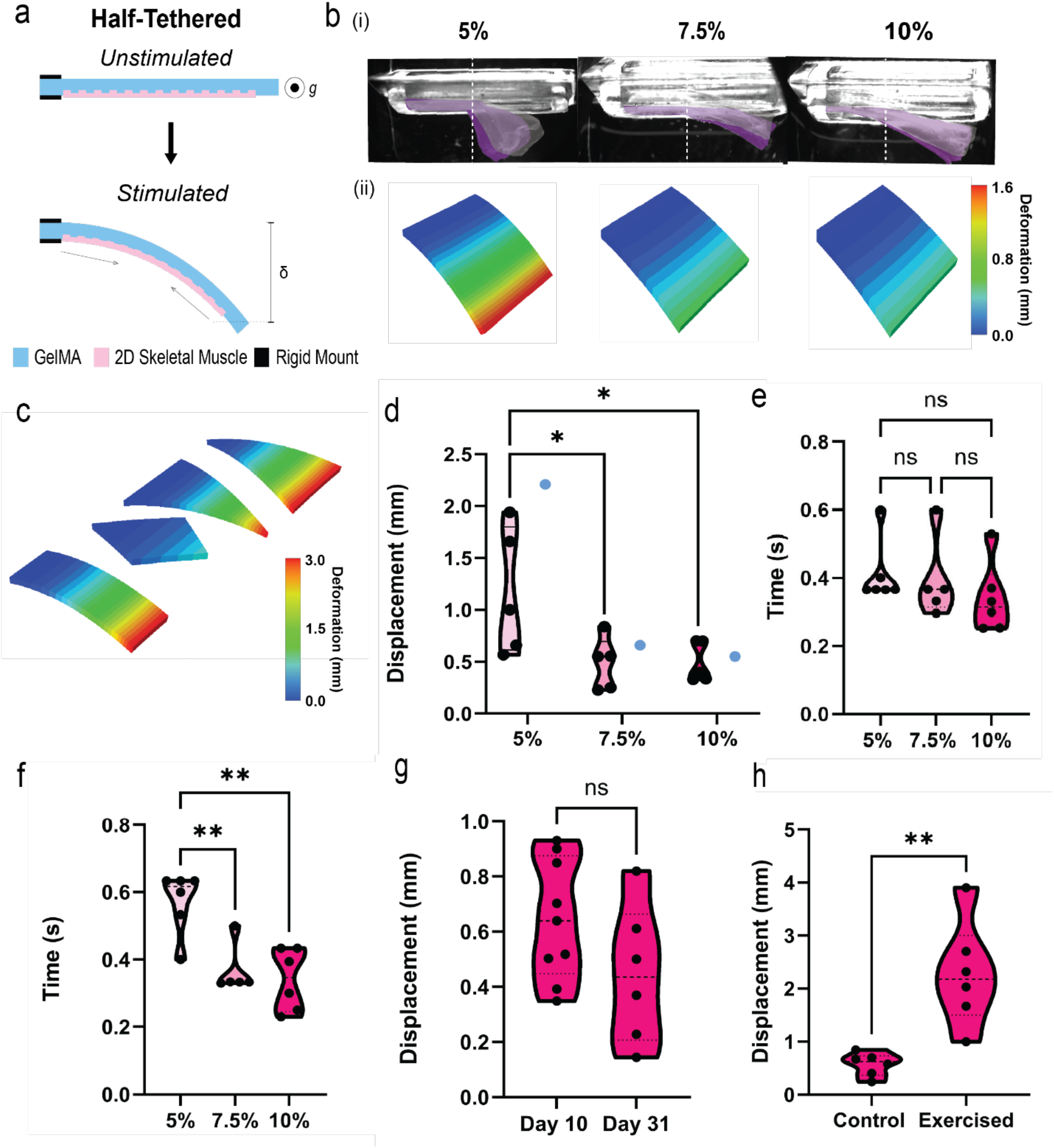
Effect of hydrogel skeleton design on out-of-plane actuation stroke. (a) Schematic of half-tethered actuation configuration. (b) (i) Images of GelMA substrates mounted on a custom jig for reproducible half-tethered actuation characterization. Muscle contraction in response to a 1Hz stimulation input (Video S1) is shown as a purple mask. (ii) Computational model predictions of peak displacements of half-tethered 5, 7.5, and 10 wt% GelMA substrates in response to muscle contraction. (c) Computational model predictions of peak displacements of different fin shapes in response to muscle contraction. (d) Empirically measured muscle stroke on half-tethered 5, 7.5, and 10 wt% GelMA substrates as compared to computational predictions (blue). Empirically measured muscle (e) contraction time and (f) relaxation time on 5, 7.5, and 10 wt% GelMA substrates. (g) Muscle stroke on 7.5 wt% GelMA substrates on day 10 and day 31 of differentiation. (h) Exercised and control muscle stroke on 7.5 wt% GelMA subsrates on day 10 of differentiation. In all panels, * and ** labels correspond to p < 0.05 and p < 0.01, respectively.

### Characterization of Untethered Actuation Performance

Efficient biohybrid robots require muscle actuators that achieve large strokes to drive functional outputs such as locomotion. Given the increased forces generated by 2D skeletal muscle thin films on GelMA substrates, we sought to characterize how these substrates impacted muscle stroke. We thus untethered actuators, while keeping one edge bound to an underlying rigid substrate, enabling robust and reproducible quantification of out-of-plane displacement in this “half-tethered” configuration (**Figure 3a**).

### Effect of Skeleton Material on 2D Skeletal Muscle Stroke

In classical mechanics of cantilever beams, the material moduli are directly and inversely proportional to the bending stiffness and force-induced deflection, respectively. Using the rheological properties determined for different GelMA concentrations, and empirically measured 2D muscle thin film properties when tethered, a finite element analysis (FEA) model was created to estimate a baseline prediction for stroke output on different substrate stiffnesses and shapes (**Figure 3b-c)**. Computational models predicted increased deformation on softer substrates, as expected, and also predicted no significant difference in deformation between rectangle, triangle, and inverted triangle shapes of normalized muscle area and parallel muscle alignment. As a result, rectangular fins of varying GelMA concentrations were tested for ease of fabrication. Empirical validation of rectangular fins with 25 µm square parallel grooves on different GelMA formulations demonstrated that the computational model accurately predicted experimental results (**Figure 3d)**. Interestingly, despite reduced force generation, the 5 wt% GelMA skeleton yielded an average stroke of 1.17 +/-0.6 mm, significantly greater than strokes of 0.48 +/0.25 mm and 0.49 +/-0.19 mm observed in 7.5 wt% and 10 wt% GelMA, respectively (**Video S1**). However, this larger stroke magnitude came at the cost of increased standard deviations in fin displacement, potentially corresponding to observed reductions in fiber alignment efficiency (**Figure 2f**), which would limit reproducibility of actuation in robotic applications. Importantly, observed strokes on all GelMA substrates were significantly higher than previously observed in 2D skeletal muscles cultured on fibrin hydrogels (∼50 µm in tissues of similar size).^[37]^

We also characterized contractile dynamics across all GelMA formulations at 1Hz stimulation (**Figure 3e and 3f)**. The time required to reach peak force, or “contraction time”, was consistent across all GelMA formulations. However, the time required to relax from peak force to baseline, or “relaxation time” was significantly lower for 7.5 wt% and 10 wt% as compared to 5 wt% GelMA. This may be explained by the higher stiffness of the GelMA substrate at higher wt% values, resulting in a quicker “spring back” effect. Since we observed no appreciable difference in the stroke magnitude, dynamics, or reproducible alignment of muscles on 7.5 wt% and 10 wt% GelMA substrates, the former was chosen for future experiments to limit material waste.

To further characterize untethered longevity of muscle thin films on 7.5 wt% GelMA substrates, actuator stroke was calculated on day 10 and day 31 of muscle differentiation, showcasing stable long-term function (**Figure 3g)**. To improve muscle performance further, we leveraged the optogenetic nature of the tissues to perform non-invasive exercise via 470 nm light stimulation, following previously optimized parameters.^[49,50]^ Specifically, we implemented a daily exercise regimen of 2 Hz optical stimulation for 15 minutes, and observed a significant 4X increase in stroke to 2.3 mm compared to a non-exercised control. Following this finding, a daily exercise routine was followed for all future experiments.

### Local Control of Multi-Limbed 2D Skeletal Muscle Thin Film Actuator

The optimized substrate topography and stiffness determined in Figures 1-3 were leveraged to fabricate a robot composed of 2 independent fins. Each fin was 7 mm wide,15 mm long, and 0.5 mm thick with a patterned muscle area of 5.8 mm width and 13 mm length (**Figure 4a**). Computational modeling predicted equal displacement for each fin in response to selective muscle actuation, as a result of the symmetric design (**Figure 4b**). The fins of fabricated robots were then actuated separately and together via spatially targeted optical stimulation at 470 nm (**Figure 4c**). We observed that with independent stimulation, the target fin contracted while the opposing fin remained stationary, matching simulation prediction of both contraction magnitude and independence. With dual fin synchronous stimulation, both fins actuated synchronously. These experiments showcased that the robot was capable of 3 independent modes of movement: left fin actuation, right fin actuation, and dual fin actuation (**Video S2**).

**Figure 4.**
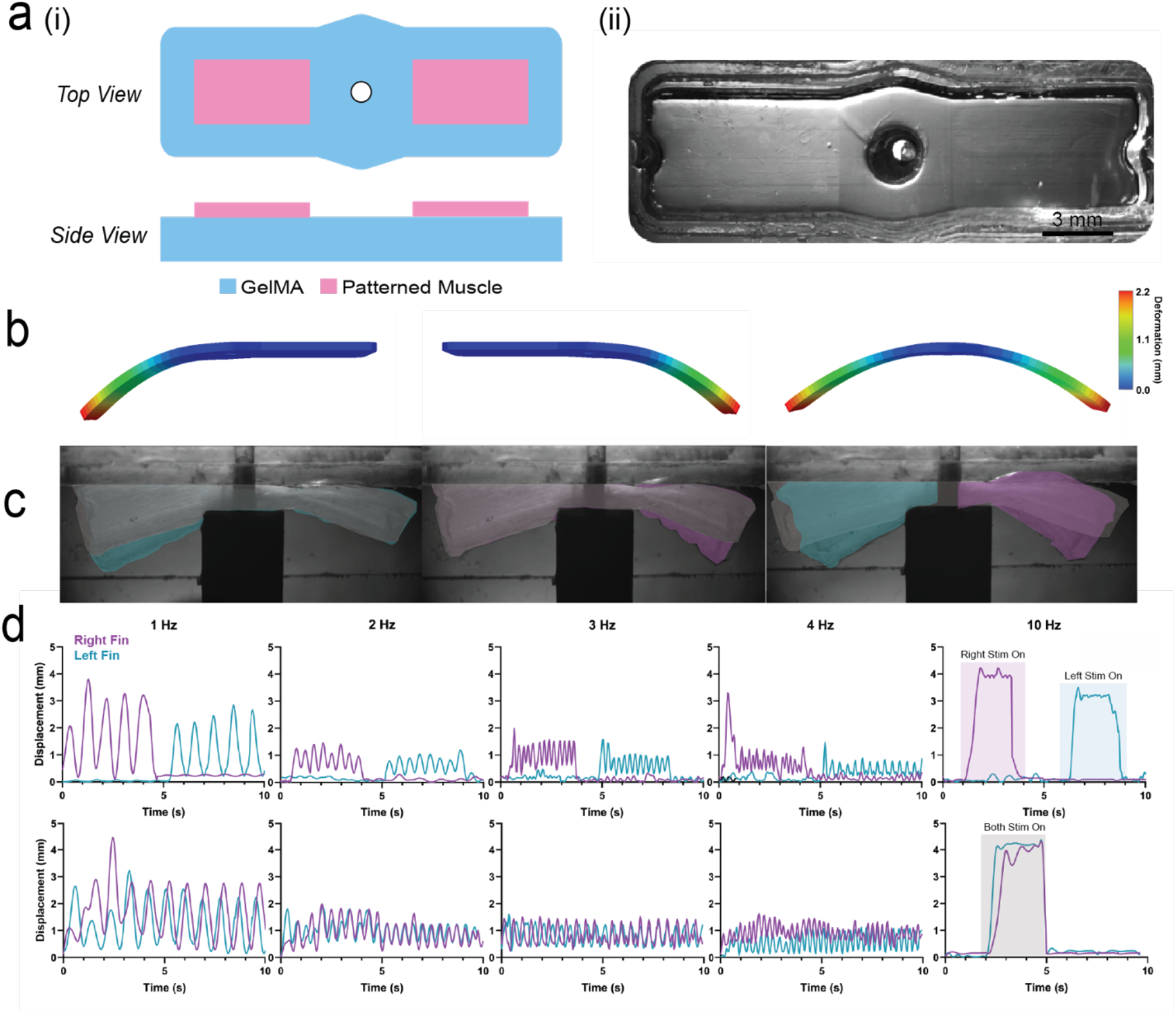
Demonstration of local control of 2-fin robot actuated by 2D skeletal muscle thin films. (a) Schematic (i) and real image (ii) of 2-fin robot. (b) Computational model predictions of left fin, right fin, and dual fin stimulation. (c) Time lapse images of 1Hz stimulation of left fin, right fin, and both fins. (d) Muscle stroke in response to left fin, right fin, and dual fin actuation at varying 1, 2, 3, 4, and 10 Hz frequencies. Tetanic contraction is observed at 10 Hz.

To characterize how the 2-fin actuator operated at different frequencies, a sweep of 1-10 Hz optical stimulation frequencies was input to the actuator and displacement was measured using a point tracking algorithm in MATLAB (**Figure 4d)**. Corresponding to both native muscle tissue and previous studies in engineered 3D skeletal muscle tissues,^[21,45,51,52]^ the dynamic range of 2D muscle contraction decreased with higher frequencies, as the muscle did not have time to fully relax before receiving the next contractile stimulus input. Also similar to 3D skeletal muscle, tetanic contraction (i.e. holding position at peak force) was observed at 10 Hz stimulation frequency, enabling sustained force output from 2D skeletal muscle thin films (**Video S3**).

### Demonstration of Multi-Directional Steering in 2D Skeletal Muscle-Powered Locomotive Robot

To establish whether 2D skeletal muscle thin films could power locomotive biohybrid robots, untethered 2-fin robots were placed in a custom maze composed of a straight section and two angled sections. After first demonstrating on/off control of locomotion (**Video S4**), the straight section was used to measure translational speed changes in the robot as a function of stimulation frequency. At 1, 2, and 3 Hz stimulation, single-fin stimulation enabled directional movement at speeds of 1.04 +/-0.38, 2.93 +/-1.5, and 3.43 +/-1.7 body lengths per minute respectively (**Figure 5a-b, Video S5**). To characterize asymmetries in fin actuation due to natural experimental variation, we compared performance between the weaker and stronger fin of each robot and observed no asymmetry in muscle stroke (**Figure S2**). We also compared directional movement in response to stimulation of the weaker and stronger fins over different frequencies. No significant changes in speed were observed between directions (**Figure 5c**), indicating that slight variations in fin fabrication did not have consequences on actuation performance or robotic function

Interestingly, a significant increase in locomotive speed was observed from 1 to 2 Hz and 1 to 3 Hz, likely due to a larger number of active contraction cycles per second. However, no significant increases in speed were observed beyond 2 Hz, potentially because of the tradeoff between reduced contractile stroke and increased number of contractions per second (**Figure 5d)**.

**Figure 5.**
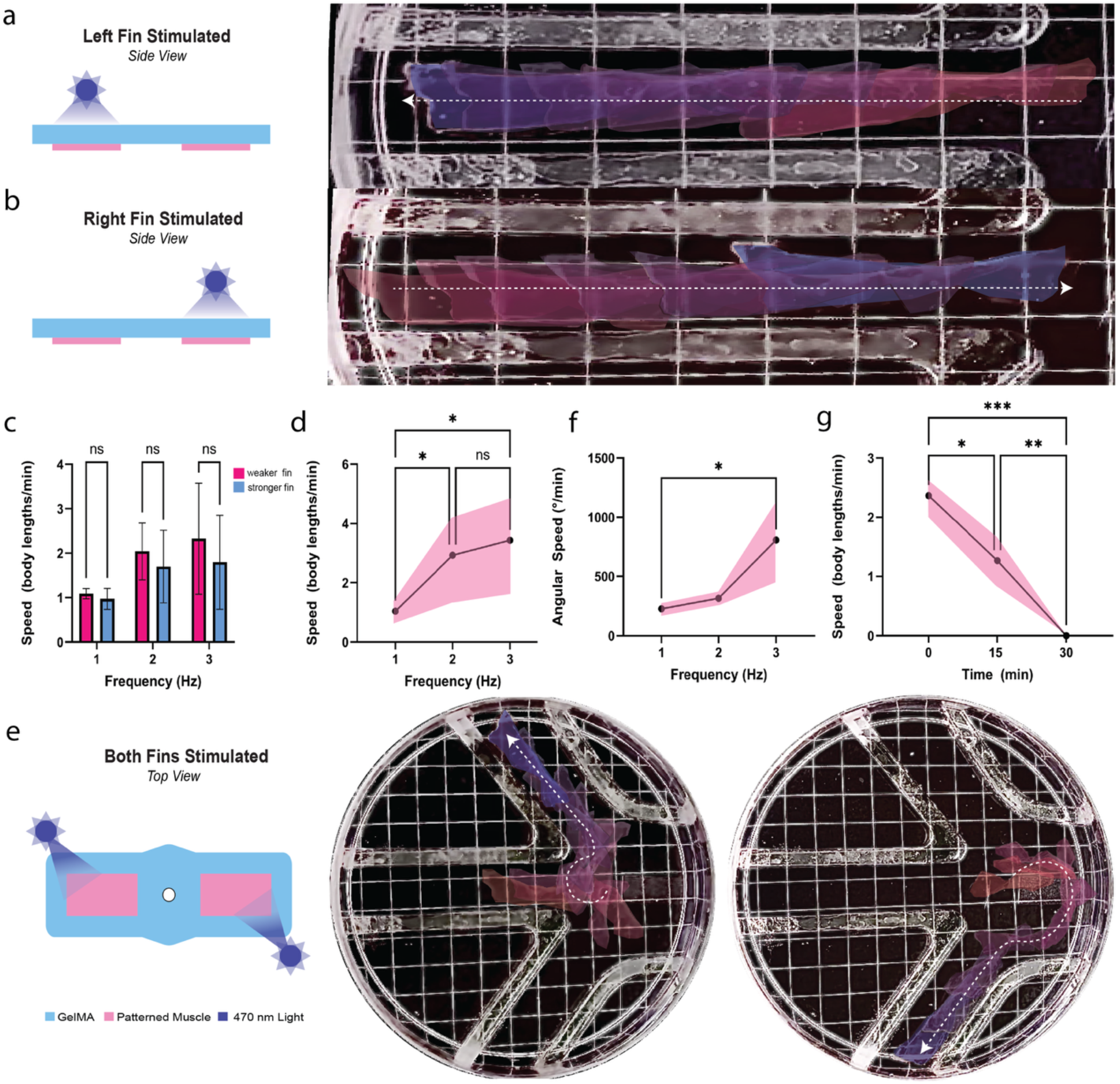
Representative demonstration of multi-directional navigation of a 2-fin robot powered by 2D skeletal muscle thin films. (a-b) Schematic of left fin and right fin stimulation (left) and time lapse images of linear motion from red to blue (right) of 2D skeletal muscle-powered robot. (c) Speed comparison of bidirectional linear speed. Since each robot had slight asymmetries in fin actuation due to natural experimental variation (**Figure S2**), we compared directional movement in response to stimulation of the faster vs slower fins over different frequencies. No significant changes in speed were observed between directions, indicating that natural variations in fin actuation did not have functional consequences on robotic function. (d) Linear speed vs. frequency of selective stimulation of left/right fins (e) Schematic of dual fin stimulation (left) and timelapse trajectory from red to blue (right) of steered robot locomotion. (f) Angular speed vs. frequency of dual fin stimulation. (g) Linear speed vs. stimulation time at 2Hz selective fin stimulation, demonstrating actuator fatigue. In all panels, *, **, and *** labels correspond to p < 0.05, p < 0.01, and p < 0.001 respectively.

Alternating stimulation of both fins enabled rotational locomotion. This turning mode was used to steer the bot into each of the separate angled tracks (**Figure 5e, Video S6**), with average rotational speeds of 226 +/-60.2, 315.5 +/-39.9, and 805+/-349 degrees/minute at 1, 2, and 3 Hz stimulation respectively (**Figure 5f, Video S7**). Once the robots were steered into the new angled path, translational motion was again engaged via 1-fin stimulation. These demonstrations highlighted controllable steering of the biohybrid robot through the maze via selective left fin, selective right fin, and dual fin stimulation.

To understand the actuation timespan of our 2D skeletal muscle thin film actuators, we performed fatigue testing at the fastest translational speed (2 Hz) via continuous optical stimulation of 1 fin for 30 minutes (**Figure 5g)**. We observed significant drops in locomotive speed from 0 to 15 minutes and from 15 to 30 minutes, with no measurable translation observed at 30 minutes. These observations are in line with fatigue studies in 3D engineered skeletal muscle actuators which also demonstrate significant drops in contractility within 30 minutes of continuous stimulation at frequencies above 1 Hz.^[51,52]^

## DISCUSSION AND CONCLUSION

Gold standard methods for fabricating skeletal muscle-powered biohybrid robots have, to date, predominantly relied on 3D muscle tissues that require large cell numbers and complex multi-step fabrication techniques while offering limited precision in muscle fiber alignment. In this work, we leverage a 1-step hydrogel micro-molding and simple cell seeding approach to fabricate 2D skeletal muscle monolayers, or thin films^[35]^. We demonstrate that tuning hydrogel stiffness and microscale topography enables precisely patterning highly contractile muscle thin films that generate mN-scale forces and mm-scale strokes from cm-scale tissues, far surpassing the performance of previous 2D skeletal muscle tissues cultured on ultra-soft hydrogels.^[46]^. Moreover, we show that, as compared to muscle monolayers cultured on ultra-soft hydrogels which cannot operate untethered for > 10 minutes, our 2D skeletal muscle thin films remain operational while untethered and in differentiation for > 30 days, corresponding to a ∼4500X increase in lifespan.

Leveraging optimized stiffness and groove parameters, we demonstrated that a symmetric 2-fin robot can be spatially controlled with light, resulting in 3 types of independent actuation modes: bidirectional linear motion with individual fin stimulation and turning with dual fin stimulation. Leveraging these 3 locomotive modes, we demonstrated multi-directional optically-guided steering through a maze. These findings can inform future biohybrid robot designs that leverage efficient use of 2D skeletal muscle thin film actuation to create multi-degree of freedom movement. While other works have shown selective stimulation and steering of skeletal muscle-powered locomotive robots, these works rely on multiple independent 3D tissues to enable multi-degree of freedom movements, and thus require significantly more cells to fabricate.^[11,21,53]^ 3D muscle-powered robots also require other components such as linkages and hinges to produce out of plane movement.^[54,55]^ These synthetic components, while functional, cannot respond to their surroundings, thus reducing the adaptive capacity of biohybrid machines. Using 2D skeletal muscle thin films enables multi-degree of freedom and out-of-plane movements, while also allowing for both simpler fabrication and greater adaptability.

Comparing our 2D skeletal muscle thin films to 3D skeletal muscle tissues generated from the same cell line (C2C12 myoblasts) in prior studies^[21,22,24–27,45,56]^ showcases that our thin films generate significantly higher isotonic contractile forces with low initial cell counts (**Figure 6a)**. Increasing mechanical output for lower fabrication input could be beneficial for high throughput and large-scale fabrication of biohybrid machines. To compare a normalized metric across this and prior studies, we also calculated force density (defined as isotonic contractile force as a function of muscle volume) for our 2D skeletal muscle thin films and previously reported 3D skeletal muscle tissues (**Figure 6b-c**). This comparison highlighted that our 2D actuators increased force density by 20X, highlighting more efficient use of muscle cells.

**Figure 6.**
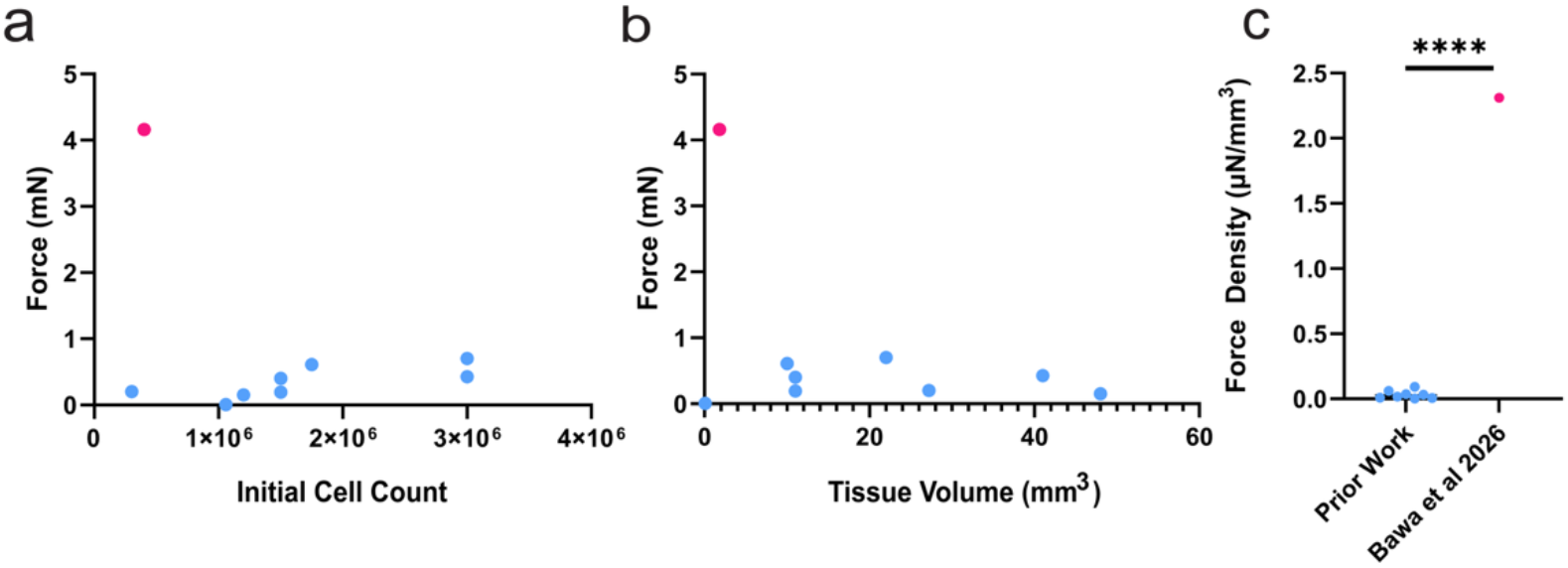
Comparison of 2D skeletal muscle thin film actuators to 3D skeletal muscle actuators. **(a)** Comparison of muscle force output vs. initial cell count used to fabricate tissues for 3D skeletal muscle actuators composed of C2C12 myoblasts^[21,22,24–27,45,56]^ (blue) and 2D skeletal muscle thin films composed of C2C12 myoblasts in this study (pink). Initial cell count was calculated using reported cell seeding densities and volumes. Isotonic contractile force values were either reported in prior studies directly, or calculated by multiplying skeleton stiffness by muscle stroke. **(b)** Comparison of muscle force output vs. tissue volume for 3D skeletal muscle actuators composed of C2C12 myoblasts^[21,22,24–27,45,56]^ (blue) and 2D skeletal muscle thin films composed of C2C12 myoblasts in this study (pink). Tissue volume was calculated by measuring length, width, and thickness of tissues in all studies. **(c)** Force density comparison of prior reported 3D skeletal muscle actuators and our 2D skeletal muscle thin films fabricated from C2C12 myoblasts, highlighted a 20X increase in efficiency. **** label corresponds to p < 0.0001.

While our demonstration of untethered steered locomotion is promising for deploying 2D skeletal muscle thin films in biohybrid robots, limitations of our study include reliance on a single cell line, uniaxial muscle alignment, and simple skeleton shapes. We anticipate that future studies that leverage our 1-step STAMPing approach to pattern more complex fin and robot geometries may yield locomotive robots that achieve higher speeds, or robots designed for different applications such as gripping or pumping. An additional limitation of our study is reliance on external stimulation, rather than onboard electronics that would enable truly untethered locomotion.

Improvements in microscale light sources and portable power sources will enable, in future, designing and deploying instrumented biohybrid robots with onboard control.^[57]^

In summary, this work establishes skeletal muscle thin films as a powerful and efficient actuation platform for biohybrid robotics. By combining precise microtopographical patterning with tunable stiffness hydrogel substrates, we demonstrate the first demonstration of multi-directional steered locomotion from a 2D skeletal muscle-powered machine. Our approach offers a scalable, cell-efficient manufacturing approach that fundamentally advances the performance and controllability of living machines, laying a strong foundation for next-generation biohybrid robots powered by skeletal muscle thin films.

## Methods

### GelMA Preparation

5, 7.5, and 10% w/v Gelatin Methacrylate (GelMA) solutions with 0.5% w/v Lithium phenyl-2,4,6-trimethylbenzoylphosphinate (LAP) (900889, Sigma-Aldrich) was produced by first reconstituting 0.5% w/v LAP in DMEM (D6429-500ML, Sigma Aldrich) at 37 °C for 30 minutes until fully dissolved. After filtering the LAP solution, 7.5% w/v Bovine 225 bloom GelMA (Cytoink) was reconstituted in the LAP solution at 37 °C until fully dissolved. The 7.5% w/v GelMA solution was stored at 4°C until further use

### STAMP Fabrication

3D nano-printed stamps were used to form the microgrooved topology on GelMA substrates. The portion of the stamps containing microgrooves were first designed in CAD (Fusion) using a linear pattern tool. Then, they were printed in UpPhoto resin with a NanoOne 2-photon printer (UpNano GmbH) with a 10x objective after slicing exported .stls with UpNano’s slicer software Think3D (UpNano GmbH). Stamps were post-processed in 3 subsequent isopropyl alcohol baths for 10 minutes each and then allowed to further photo-cure for at least 12 hours before assembly to holders printed using stereolithography. Stamp holders were designed in CAD to include a hardstop against molds to leave a 500 µm gap, the desired thickness of the stamped substrate. The final designs were then sliced with Preform slicing software (Formlabs) and printed using a Formlabs 3B Stereolithography printer with Biomed Clear Resin (Formlabs). The holders were post-processed in isopropyl alcohol in the Form Wash for 30 minutes and then cured in the Form Cure for 60 minutes at 60 °C. The holders were then run through a Tuttnauer benchtop line autoclave (Tuttnauer USA Co Ltd, 2840ELD) at 121 °C using the preset plastic cycle and allowed to cool to room temperature. Subsequently, the nano-printed stamps were adhered to the holders with a thin layer of polyurethane tape.

### Fabrication of Microgrooved GelMA and Fibrin Substrates

Once the GelMA solution was prepared, it was cast in a custom designed mold, made with either a glass or acrylic base with a laser cut frame adhered on top with a laser cut polyurethane tape. The assembled molds and stamps were then sterilized via immersion in 3% H_2_O_2_ (FisherScientific) overnight at room temperature. Residual H_2_O_2_ was rinsed off all components by immersion in a PBS bath for at least 30 minutes at room temperature. Stamps were then coated with 5% bovine serum albumin (BSA) (Thermo Fisher) for at least 1 hour at 37°C. GelMA was prepared for micropatterning by warming it to 37°C. All molds were plasma treated for 30 seconds per well using a handheld oxygen plasma treater to promote forming a flat layer of gel. Once warmed, 150µL GelMA solution was placed in each mold for the single limb actuators and 300µL for the dual-limb actuators. The molded gels plus stamps were then placed over a 475nm wavelength lamp (Thorlabs) for cross-linking. The lamp was placed at 6 mW power and the gels were cross-linked for 30 seconds. Then the lamp was turned off, and the gel was left at room temperature for 10 minutes to undergo physical cross-linking. Stamps were then removed and the gels underwent a final round of crosslinking at 20mW power for 2 minutes.

To create microgrooves on Fibrin substrates, 150 µL of 8 mg/mL fibrinogen and 100 U/mL thrombin solution was prepared and cast into molds. Stamps were placed on top of the unpolymerized fibrin gels cured at 37°C for 1 hour. Stamps were then removed. All individual molds were placed in wells of a glass-bottom 6 well plate (Cellvis).

To ensure stamps could be reused with high feature fidelity, a three-part cleaning step was followed. First, stamps were soaked in a 0.1 mg/mL Liberase solution (Sigma Aldrich) for at least 1 hour at 37°C to degrade any residual GelMA stuck to the small features of the stamp. Then, stamps were re-sterilized and cleaned using H_2_O_2_ for 30 minutes at room temperature, followed by a final PBS soak for an additional 30 minutes at room temperature. After cleaning, stamps could be coated with BSA, enabling reuse.

### Myoblast Seeding and differentiation on GelMA Substrates

C2C12 myoblasts were initially cultured on plastic tissue culture treated flasks using muscle growth medium which contained 10% fetal bovine serum (A5670701, Gibco), 1% penicillin streptomycin (MT30002CI, Fisher Scientific), and 1% L-glutamine (MT25005CI, Fisher Scientific) in a high glucose formulation of DMEM with 4.5 g/L of glucose and sodium pyruvate (MT10013CV, Fisher Scientific). Myoblasts were passaged prior to reaching 70% confluence. C2C12s were prepared at a concentration of 4×10^6^ cells/mL and seeded at a cell density of 80K cells/mm^2^. Samples were incubated for 30 minutes to ensure cell attachment to the hydrogel substrates and then 3mL of culture media was added to each well. Myoblasts were maintained in molds in growth medium containing 1 mg/mL of aminocaproic acid (A2504, Sigma Aldrich). Upon reaching confluence in the substrates, myoblasts were switched to differentiation medium to induce myoblast fusion and maturation. Muscle differentiation media consisted of 10% horse serum (26050-088, Gibco), 1% penicillin-streptomycin, 1% Lglutamine, and 50 ng/mL of human insulin-like growth factor-1(I1146, Sigma-Aldrich) in a high glucose formulation of DMEM. Media was changed daily throughout the course of the experiments

### Evaluation of Muscle Shear Displacement and Force

Videos of muscle contraction were recorded in brightfield with a 10x objective at 30 fps with a Zeiss Primovert Microscope. A commercial electrical stimulation setup, C-Pace (C-Pace EM100 from ionoptix) was used for stimulation. The C-Pace Lid with electrodes was placed on top of the wells, and a 10 Vpp, 10% duty cycle, 1Hz square pulse signal was used for stimulation. After videos were collected, they were analyzed using a point tracking algorithm and an average displacement over all tracking points was determined for each video. To calculate the shear force applied by the muscle, the average displacement was divided by the thickness of the gel to determine shear strain. Using experimentally determined shear moduli for each gel type, the stress was calculated.

### Evaluation of Muscle Alignment

On day 10 of muscle differentiation, all samples were imaged on a Nikon confocal microscope at 4x. Large stitch, z-stack fluorescent images were taken to visualize the entire muscle layer. Images were then fed into a custom alignment pipeline in python, which rotated all images so that the long edge of each mold was at 90 degrees and cropped all areas outside the mold. Finally, fast Fourier transform with 180 bins was done in Fiji/ImageJ to determine a gaussian distribution (peak angle and standard deviation) of light intensity as a function of angle.

### Evaluation of Muscle Stroke

To determine stroke output of each sample, each sample was untethered from the mold. Then, using a laser cut slot on an acrylic base, each actuator was turned 90 degrees and inserted into the slot to achieve a “half-tethered” configuration. To demonstrate selective fin stimulation of the 2-fin robots, all samples were untethered and inserted into a custom jig to enable reproducible visualization of actuation. Optical stimulation of each fin at 470 nm was performed using an optical fiber with a collimating adaptor (ThorLabs). Since recorded displacements could be seen with the naked eye, a 1x magnification was used on a stereoscope (Zeiss) to visualize to record stroke displacements. Videos were fed through a KLT point tracking algorithm to calculate the peak displacement at the tip of each actuator.

### Computational Model for Material Stiffness and Fin Shape

To simulate muscle actuation on GelMA substrates, a 3D model of the gel substrate was created in Abaqus (**Figures 3c-d)**. To test different fin shapes, total muscle area and thickness and alignment were kept constant and only shape was modulated by controlling the dimensions of each shape. Experimentally determined shear moduli were entered for each gel type: 5,7.5,10% GelMA, and Fibrin. The muscle layer was set up as a thermal material that would exhibit contractile strain under a temperature drop, as previously described.^[37]^ A 3D mesh with hybrid elements was created with a mesh size of 0.3 for all simulations, and minimum step size of 0.00001 and a max step size of 0.001. Boundary conditions for single limb simulations included a fixed face at the far edge of each sample to emulate a cantilever beam. For dual limb simulations, only the middle section of the gel body was fixed.

### Rheology

Rheological characterization was performed using a TA Instrument Discovery HR-2 Hybrid rheometer equipped with an Advanced Peltier Plate temperature-control system and an 8 mm parallel-plate geometry. All measurements were conducted at 37 °C using a fixed gap height of 1 mm. Hydrogel samples were prepared in PDMS molds following the same formulation and crosslinking protocol used for the cell culture experiments. Prior to testing, samples were incubated in cell culture medium at 37 °C for 1 hour to ensure hydration and thermal equilibration. During measurements, a solvent trap was used to minimize dehydration. Amplitude sweep tests were first performed at 1 Hz over a strain range of 0.01–100% to identify the linear viscoelastic region. Based on these results, frequency sweep measurements were conducted within the linear viscoelastic regime over an angular frequency range of 0.01–100 rad. s^−1^ using strain amplitudes of 0.5% and 0.7%. Storage and loss modulus were recorded as functions of angular frequency.

### Statistical Analysis

All statistical tests were performed on GraphPad using either a One-way Anova or unpaired t-test for statistics with only two groups, with *, **, ***, and **** labels corresponding to p < 0.05, p < 0.01, p < 0.001, and p < 0.0001 respectively.

## Funding Acknowledgement

This work was funded by the DoD Office of Naval Research Young Investigator Program (N00014-24-1-2060, awarded to R.R.), the DoD DURIP Program (W911NF-24-1-0106, awarded to R.R.), the National Science Foundation Graduate Research Fellowship Program (M.B.), and the MIT Engineering Excellence Postdoctoral Fellowship Program (A.B.). Our research was partly conducted within the Koch Institute’s Robert A. Swanson (1969) Biotechnology Center, supported by a core grant P30-CA14051 from the NIH National Cancer Institute, and benefited from technical support from the Peterson (1957) Nanotechnology Materials Core Facility.

