## Supplemental Information for "2D Skeletal Muscle Thin Film Actuators Enhance Efficiency of Biohybrid Robots"

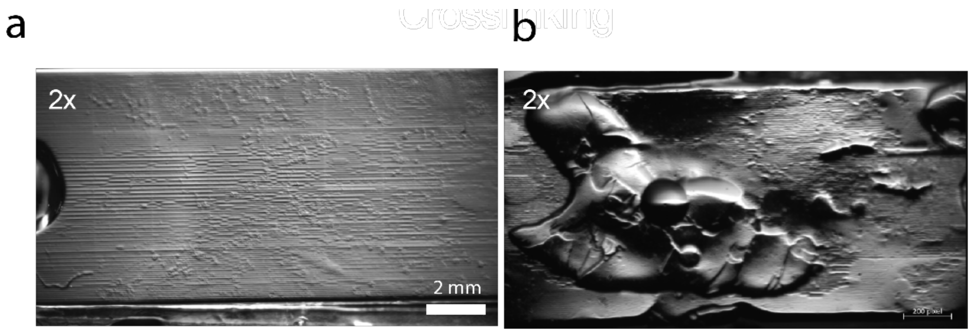


**Figure S1.** Comparison of STAMPing GelMA hydrogels with **(a)** and without **(b)** physical crosslinking.

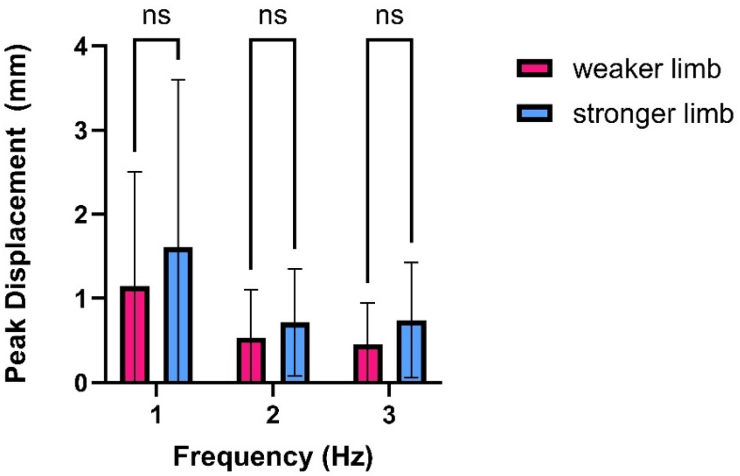


**Figure S2.** Comparison of the weaker limb (limb with lower peak displacement) and stronger limb during half-tethered frequency sweep tests of 2-fin actuators.

**Link to Supplemental Videos**
